# Evidence accumulation during perceptual decision-making is sensitive to the dynamics of attentional selection

**DOI:** 10.1101/537910

**Authors:** Dragan Rangelov, Jason B. Mattingley

**Affiliations:** Queensland Brain Institute, The University of Queensland, St Lucia, QLD 4072, Australia; School of Psychology, The University of Queensland, St Lucia, QLD 4072, Australia

**Keywords:** decision making, visual attention, encephalography, mixture distribution modelling, frequency tagging, multivariate analyses

## Abstract

The ability to select and combine multiple sensory inputs in support of accurate decisions is a hallmark of adaptive behaviour. Attentional selection is often needed to prioritize stimuli that are task-relevant and to attenuate potentially distracting sources of sensory information. As most studies of perceptual decision-making to date have made use of task-relevant stimuli only, relatively little is known about how attention modulates decision making. To address this issue, we developed a novel ‘integrated’ decision-making task, in which participants judged the average direction of successive target motion signals while ignoring concurrent and spatially overlapping distractor motion signals. In two experiments that varied the role of attentional selection, we used linear regression to quantify the influence of target and distractor stimuli on behaviour. Using electroencephalography, we characterised the neural correlates of decision making, attentional selection and feature-specific responses to target and distractor signals. While targets strongly influenced perceptual decisions and associated neural activity, we also found that concurrent and spatially coincident distractors exerted a measurable bias on both behaviour and brain activity. Our findings suggest that attention operates as a real-time but imperfect filter during perceptual decision-making by dynamically modulating the contributions of task-relevant and irrelevant sensory inputs.

## Introduction

Cognition can be conceived of as a cascade of processes, beginning with the encoding of sensory input and ending with appropriate behavioural choices. Sequential-sampling decision-making models have had considerable success in describing choice behaviour in terms of both behavioural performance and neural activity (Forstmann, Ratcliff, & Wagenmakers, 2016; O’Connell, Shadlen, Wong-Lin, & Kelly, 2018; Ratcliff, Smith, Brown, & McKoon, 2016). According to these models, decision-making relies on a central decision variable, an abstract quantity that accumulates in time toward one of several possible choice alternatives. On these accounts, a choice is made once the decision variable reaches threshold. In one widely employed decision-making task – the random-dot motion discrimination task (Ratcliff et al., 2016; Shadlen & Kiani, 2013) – participants are shown fields of randomly moving dots and are asked to judge the motion direction of a fraction of the dots that move coherently. Studies in both animals (Shadlen & Kiani, 2013) and humans (Heekeren, Marrett, & Ungerleider, 2008; Kelly & O’Connell, 2013; Liu & Pleskac, 2011; O’Connell, Dockree, & Kelly, 2012; Ratcliff, Philiastides, & Sajda, 2009; Twomey, Kelly, & O’Connell, 2016) have shown that neural activity recorded during random-dot motion discrimination follows a trajectory that resembles the accumulation of a decision variable, thereby lending neurobiological support to sequential-sampling decision-making models.

To date, much has been learned about simple perceptual decision-making in humans and other animal species (Hanks & Summerfield, 2017; O’Connell et al., 2018; Summerfield & Tsetsos, 2015). In the real world, however, decisions are often more complex, requiring judgements on stimuli that are separated in time and space, and which may or may not be task relevant. For example, deciding to cross a busy road requires us to integrate the motion directions of vehicles coming from both the left and right sides, while disregarding any movements made by people or objects on the sidewalk. In recent years there has been increased interest in studying behaviour and neural activity in more complex decision-making tasks (Churchland & Ditterich, 2012; Churchland, Kiani, & Shadlen, 2008; Itthipuripat, Cha, Deering, Salazar, & Serences, 2018; Kohl, Spieser, Forster, Bestmann, & Yarrow, 2018; Leite & Ratcliff, 2010; Spitzer, Blankenburg, & Summerfield, 2016; Spitzer, Waschke, & Summerfield, 2017; Wyart, Myers, & Summerfield, 2015). To increase decision complexity, many of these studies have varied the number of response alternatives (e.g., two vs. four), and have found significant costs associated with a larger number of potential responses (e.g., Churchland et al., 2008; Leite and Ratcliff, 2010; Itthipuripat et al., 2018; Kohl et al., 2018). To date, however, the question about how the brain regulates decision-making when it receives competing – task-relevant and irrelevant – sensory inputs has received little attention in the literature. The central aim of the present study was to characterise the effects of selective attention on perceptual decision-making. Specifically, we were interested in measuring the degree to which relevant and irrelevant signals are integrated into a decision variable that is used as the basis for behavioural choices.

According to sequential-sampling decision-making models, selective attention should mediate sensory input and evidence accumulation by filtering out task-irrelevant inputs and preventing them from being accumulated into the decision variable (Leite & Ratcliff, 2010; Smith & Ratcliff, 2009; White, Ratcliff, & Starns, 2011; Wyart et al., 2015). This framework raises at least two inter-related questions. First, what is the relationship between the time-course of attentional selection and evidence accumulation, and second, how effective is attention in mediating sensory encoding and evidence accumulation? Conceptually, selection and accumulation could take place sequentially so that only prioritised, task-relevant input is accumulated. Alternatively, the two processes could take place in parallel, so that task-irrelevant inputs might also impact upon evidence accumulation to some extent, particularly in the early stages of attentional selection.

Studies on visual attention have repeatedly shown robust behavioural and neural costs when task-irrelevant, distracting stimuli are presented together with task-relevant target stimuli (e.g., S. K. Andersen & Müller, 2010; Eriksen, 1995; Theeuwes, 1992). While the literature agrees that attentional selectivity is imperfect, relatively little is known about the effects of attention on the dynamics of evidence accumulation. When both target and distractor signals are present, attention might influence decision making in at least two ways. On the one hand, target and distractor stimuli could each trigger a separate decision-making process, in which case attention would modulate the probability with which the two processes drive behaviour. On the other hand, if attention operates as a filter between sensory input and evidence accumulation, it would be expected to modulate the degree to which target and distractor signals are accumulated into a single decision variable. The critical distinction between these two accounts relates to the content of decision variable. In the former case, the decision variable could reflect either the target or distractor signal, whereas in the latter it would be a mixture of *both* target and distractor signals.

In the present study, across two experiments, we used random-dot displays comprising ***two*** simultaneous, intermingled fields of dots, one in a cued (target) colour, the other in an uncued (distractor) colour (see Fig. 1). Past research has suggested that the number of targets may influence the effectiveness of attentional selection (Frătescu, Van Moorselaar, & Mathôt, 2019; Hollingworth & Beck, 2016; van Moorselaar, Theeuwes, & Olivers, 2014). Motivated by these findings, we wanted test whether and to which extent the complexity of attentional set influences evidence accumulation. To that purpose, in different blocks of trials, we cued our participants to monitor for either a single target colour (e.g., blue) or two target colours (e.g., blue and yellow). In each trial, participants were presented with successive epochs of dot motion (each containing the two overlapping dot fields, one in the target colour and the other in a distractor colour). At the end of the trial, participants’ were instructed to reproduce the ***average target motion direction***, ignoring the directions of the concurrent distractor-dot fields. Thus, for example, if target motion direction was 90° for the first display and 110° for the second, the *average* motion direction would be 100°. Participants made their responses on a continuous scale by rotating a dial to match their judgement of the average motion direction of the two targets. This approach enabled us to characterise the precision with which participants integrated the two target signals and to separate noisy target responses from random guesses using mixture distribution modelling (Schneegans & Bays, 2016). Finally, a continuous nature of our response variable enabled us to use linear regression to quantify the degree to which presented target and distractor motion signals influenced the responses.

In addition to these behavioural measures, we recorded participants’ neural activity using electroencephalography (EEG). We focused in particular on the temporal dynamics of decision processes as reflected in the central-parietal positivity (CPP). Previous research has shown that only task-relevant stimuli evoke the CPP (Loughnane et al., 2016) and that the CPP is observed even when no response is required (Twomey et al., 2016), suggesting that this component is neither a passive sensory response nor a correlate of motor planning. Further, the CPP time-course closely resembles the time-course of evidence accumulation: specifically, its amplitude builds up gradually (O’Connell et al., 2012) and the build-up slope is proportional to stimulus quality (Kelly & O’Connell, 2013) suggesting that the CPP is intimately related to decision making.

**Figure 1.**
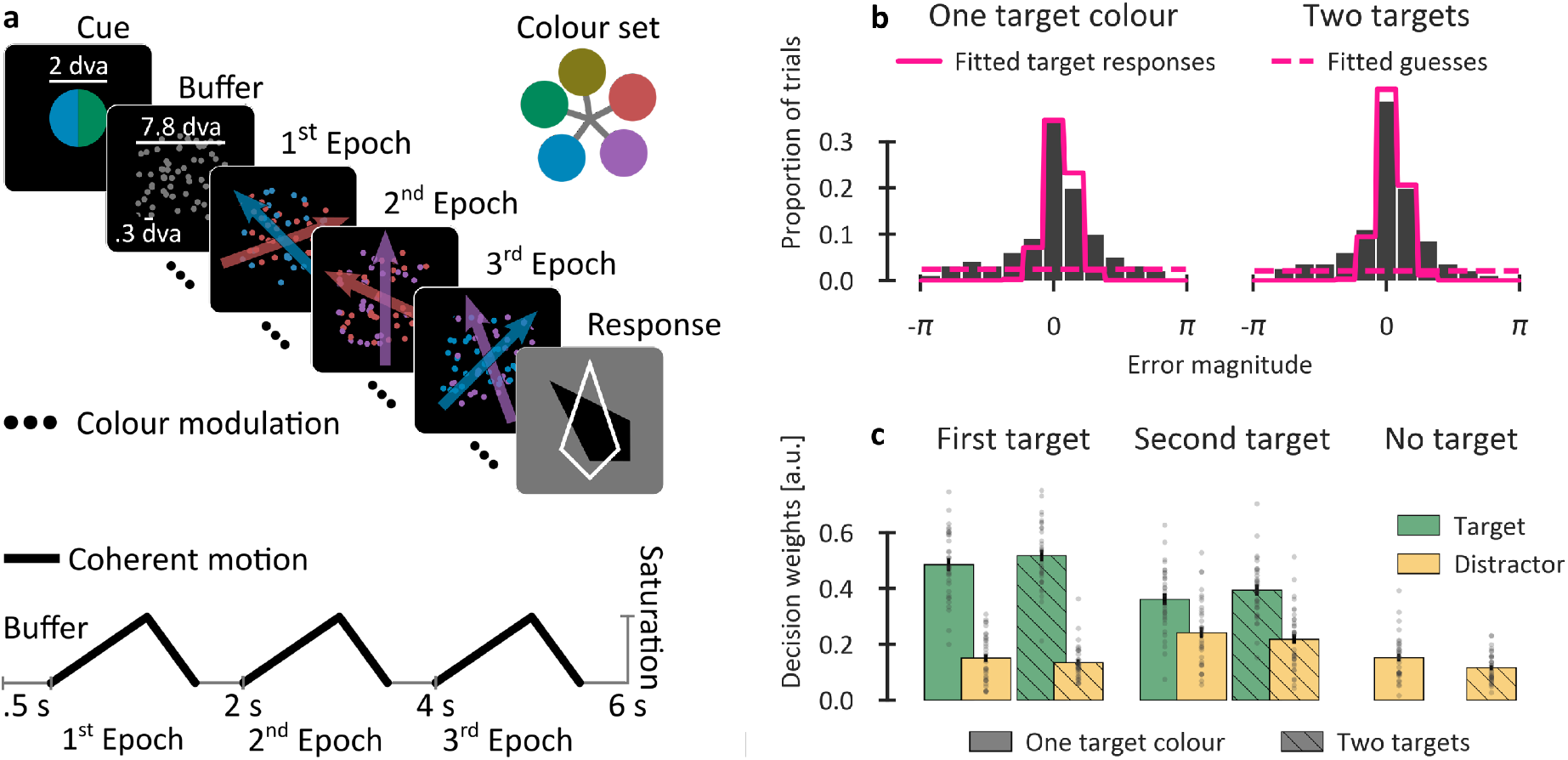
Schematic of the stimuli, task and the behavioural results in Experiment 1. **a,** Example stimulus sequence from a typical trial (above) together with the timing of colour saturation modulation and coherent-motion onsets (below). Participants had to reproduce the *average* motion direction of target-coloured dots by adjusting the orientation of a response dial (black pointer). After indicating their response they received feedback on accuracy within the same display (white pointer). **b**, Group-level histograms (bars) of the error magnitude (reproduced motion direction minus expected). The pink lines show fitted distributions of target responses and guesses using mixture-distribution modelling of each individual participant’s data. **c**, Average decision weights showing the contributions of each presented motion direction to the decision. Since there were no significant differences (t < 1) between the two distractor motions in the “No target” condition, only the average of the two is shown. Grey dots denote individual participants.

## Experiment 1

### Materials and Methods

#### Participants

36 healthy volunteers (19 females, average age 21.2 **±** 2.5 years) participated in Experiment 1. All had normal or corrected-to-normal visual acuity and normal colour vision. Three participants were identified as outliers based on their behavioural performance and/or EEG signals (see below for details), leaving 33 participants in the final sample. The study was approved by the Human Research Ethics Committee of The University of Queensland (Approval 2016001247), and was conducted in accordance with the Human Subjects Guidelines of the Declaration of Helsinki. Participants provided written informed consent prior to taking part in the study.

#### Stimuli, task and procedure

At the beginning of each trial of Experiment 1, a central disk, shown for 1 s, cued participants to the target colour (Fig. 1a). In separate blocks of trials, participants were cued to look for either one target colour (e.g., blue), or two different target colours (e.g., blue and yellow). The cue was followed by a .5 s buffer period showing a field of 80 grey, randomly-moving dots. Then, over a 1.5 s period, the dots’ colour saturation gradually increased before returning to grey. The increase in saturation revealed two overlapping, coloured-dot patches moving in different directions. Each trial comprised three epochs of colour modulation – in two of these epochs, one dot patch was rendered in the target colour, and the other appeared in a distractor colour. In the remaining (“no target”) epoch, neither of the patches had a target colour. Target patches were displayed equally often in the first, second and third epochs of a trial. Participants were asked to monitor successively presented epochs and to indicate the ***average direction*** of target-motion events at the end of the trial using a response dial. A response feedback symbol showing the correct answer was presented after every trial. Participants were instructed to prioritise response accuracy over response speed, and the response times were not analysed further. Participants completed 30 practice trials followed by six blocks of 44 trials (264 trials in total) which took approximately one hour.

In Experiment 1, dot-motion was generated at a suprathreshold level (40% coherence), with a speed of 19°/s, 100 ms dot-life, and with different signal-dots across frames and random-walk noise-dots (Scase, Braddick, & Raymond, 1996). Although coherent motion was present throughout the epoch, different motion directions were discernible only at higher colour saturations. The motion directions were uniformly sampled from a 0°–359° range in 1° steps, with the target-distractor and target-target difference remaining within ±30°–150°. Five easily distinguishable colours were selected in the HSL colour-space with a minimum hue difference of 72° and medium saturation and brightness (both 50). Two randomly-selected colours served as targets and two as distractors. In two target-colour trials, there was an equal number of trials in which the target colour repeated or changed between two target-present epochs within the trial.

#### Apparatus

The experiments were conducted in a dark, acoustically and electromagnetically shielded room. The stimuli were presented on a 24” monitor with 1920×1080 resolution and a refresh rate of 144 Hz. The experimental software was custom-coded in Python using the PsychoPy toolbox (Peirce, 2007, 2009). EEG signals were recorded using 64 Ag-AgCl electrodes (BioSemi ActiveTwo) arranged in the 10-20 layout, and sampled at 1,024 Hz.

#### Behavioural analyses

To identify outlier participants, the distributions of error magnitudes (i.e., the angular difference between the response and the correct answer) were compared to a uniform distribution (i.e., pure guessing) using the Kolmogorov-Smirnov test. Participants for whom the probability of the null hypothesis (i.e., a uniform distribution of error magnitudes) exceeded .001 were removed from further analyses.

The remaining distributions per experimental condition and per participant were fitted to a theoretical model (Schneegans & Bays, 2016), and responses were separated into noisy target responses and random guesses. We estimated two parameters per participant and experimental condition: (i) the response precision of the target responses (K parameter) and (ii) the guessing rates (a).

Finally, we quantified the degree to which presented motion signals influenced the response. To that purpose, a multiple linear regression (OLS) model with a term for each of the presented motion directions, expressed as complex numbers, was fitted to the responses, separately per participant and experimental condition. Transforming the angles into complex numbers was necessary, as both our independent and the dependent variables were circular variables. We could not use real values, because real-valued linear regression would treat the 0° and 360° angles as different even though the two are in fact alternative representations of the same angle. The absolute value of the resulting regression coefficients reflects the influence of each of the presented coherent motion signals on the response, i.e., its *decision weight*. As the decision weights were always positive, we used permutation testing (10,000 permutations) to generate the empirical null distribution against which the empirical decision weights were tested.

#### EEG analyses

EEG signals were analysed using the MNE-Python toolbox (Gramfort et al., 2013). The data were re-referenced offline to the average electrode, low-pass filtered at 99 Hz and notch-filtered at 50 Hz to eliminate line noise. The recorded signal was pre-processed using the FASTER algorithm for automated artefact rejection (Nolan, Whelan, & Reilly, 2010). The pre-processed signal was down-sampled to 256 Hz, segmented into epochs between the cue-onset and the response-display onset, baseline-corrected relative to -.1–0 s pre-trial and linearly de-trended. Outlier epochs and participants were identified using the FASTER algorithm and removed from further analyses.

The CPP was computed by averaging Cz/CPz electrodes across trials. Each epoch within a trial was extracted from the onset of the increase in colour saturation and baseline-corrected relative to a common baseline during -.1–0 s, averaged across all epochs. We quantified the CPP onset latency, slope and peak amplitude. The peak CPP amplitudes were identified using the jackknife-method (Ulrich & Miller, 2001). Using the same method, the CPP onset latencies were computed at 50% peak amplitude. The CPP slopes were computed as the amplitude difference between -.05 s and .05 s relative to the ERP onset, and divided by .1 s so as to yield μV/s.

To characterise feature-specific neural responses to coherent motion, we used a multivariate analysis approach described previously (Ede, Chekroud, Stokes, & Nobre, 2018, 2019; Wolff, Jochim, Akyürek, & Stokes, 2017). This analysis splits the whole data set into training and test epochs and uses the results derived from the training set to characterise the brain’s response in the test epochs. We used an iterative, leave-one-out algorithm in which each epoch of coherent motion served as a test epoch and the remaining epochs served as the training epochs. Two separate analyses were conducted, one for the target motion direction, the other for the distractor motion direction. The training epochs were first sorted into 16 equidistant bins (*β* centres range = −180–180°, width = 90°) based on the angular distance between motion directions in the test and the training epochs. The epochs within a bin were then averaged, yielding a bin-specific ERP. For each time sample, a multidimensional Mahalanobis distance between the test and each bin ERP was computed using temporally smoothed (Gaussian window, SD = 16 ms) and spatially filtered (surface-Laplacian) signals from all 64 electrodes. This analysis yielded a distance matrix (*D*_*M bins × n samples*_) containing the distance (*d_mn_*) between a test epoch and a given bin *m* for time sample *n*. The distances were then normalised by subtracting the mean bin distance per time sample 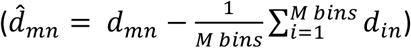, temporally smoothed using a Gaussian window (SD = 16 ms), and expressed as a similarity matrix by inverting the sign 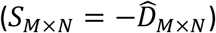.

If the test epoch contains feature-specific signals, the similarity matrix should contain the largest values for bins with the smallest angular difference relative to the test direction (e.g., ±15°), and the smallest values for bins with the largest difference (e.g., ±180°). In a final step, we computed an aggregate measure of motion-specific response (i.e., motion tuning) by convolving the similarity matrix with a vector of cosine-transformed bin centres (*Tuning_N_* = cos (*β_M_*) · *S_M×N_*). This non-parametric measure provides a robust index of feature-specific brain responses. It returns a value of zero when the similarity is the same across all bins and increases gradually with sharpening of the similarity profile across bins.

### Results

To quantify how well participants could perform the task, we used mixture distribution modelling to estimate the response precision (K) and guessing rates (a) per participant and experimental condition. Initial analyses showed no differences between repetitions and changes of target colour within a trial (all ts < 1), so these two trial types were averaged for further analyses. The average response precision was high both in the one-and two-target colour conditions (K_M/SEM_ = 10.52/1.70 and 11.79/2.08), and did not differ significantly between conditions (F < 1). Overall, the full-width-at-half-maximum (FWHM) of the fitted distribution of target responses was around 40°. For reference, the average tuning width of individual neurons in macaque area MT is about 60° FWHM (Treue, Hol, & Rauber, 2000). The average guessing rates (a) were around 26%, and did not change across experimental conditions (F < 1).

To quantify how much each of the presented target and distractor signals influenced observers’ average-motion decisions, we used linear regression modelling with complex-valued data (see Methods). The decision weights for all motion signals in all experimental conditions were significantly different from zero (all p < .001), indicating that both task-relevant and task-irrelevant signals contributed to the decision. As with the mixture distribution modelling, initial analyses of decision weights revealed no significant differences between target colour repetitions and changes (all Fs < 1.97, all ps > .170), so the two conditions were averaged for further analyses. Overall, the weights for targets (M/SEM = .44/.01, Fig. 1c) were much higher (F_2,64_ = 326.80, p < .001, 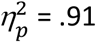) than the weights for either concurrently presented distractors (.19/.01) or distractors presented on their own (.13/.01). Unexpectedly, there was also a reliable order effect, with higher target weights for the first epoch than for the second (.50/.01 vs. .38/.02, t32 = 4.74, p < .001); analogously, distractor weights were lower for the first epoch than for the second (.14/.01 vs. .23/.01, t_32_ = 4.68, p < .001). Overall, the decision weight analyses revealed that distractors, presented both with and without the target, contributed significantly to the response, albeit to a lower degree than target motions.

We next characterised the temporal dynamics the CPP ERP. We were particularly interested in the CPP in no-target epochs: if attentional selection and evidence accumulation operate sequentially, then attention should suppress motion signals, predicting no CPP in this epoch. By contrast, if selection and accumulation take place concurrently, there should be a significant CPP even in the no-target epoch. Inspection of the ERP topographies (Fig. 2a) revealed a central-medial positivity consistent with the usual CPP component (O’Connell et al., 2012). The CPP (Fig. 2b) closely followed the time-course of colour modulation, with three distinct peaks coinciding with the three epochs within each trial. Between epochs, as the colour saturation of the dots returned to grey, the CPP amplitude also returned close to baseline.

**Figure 2.**
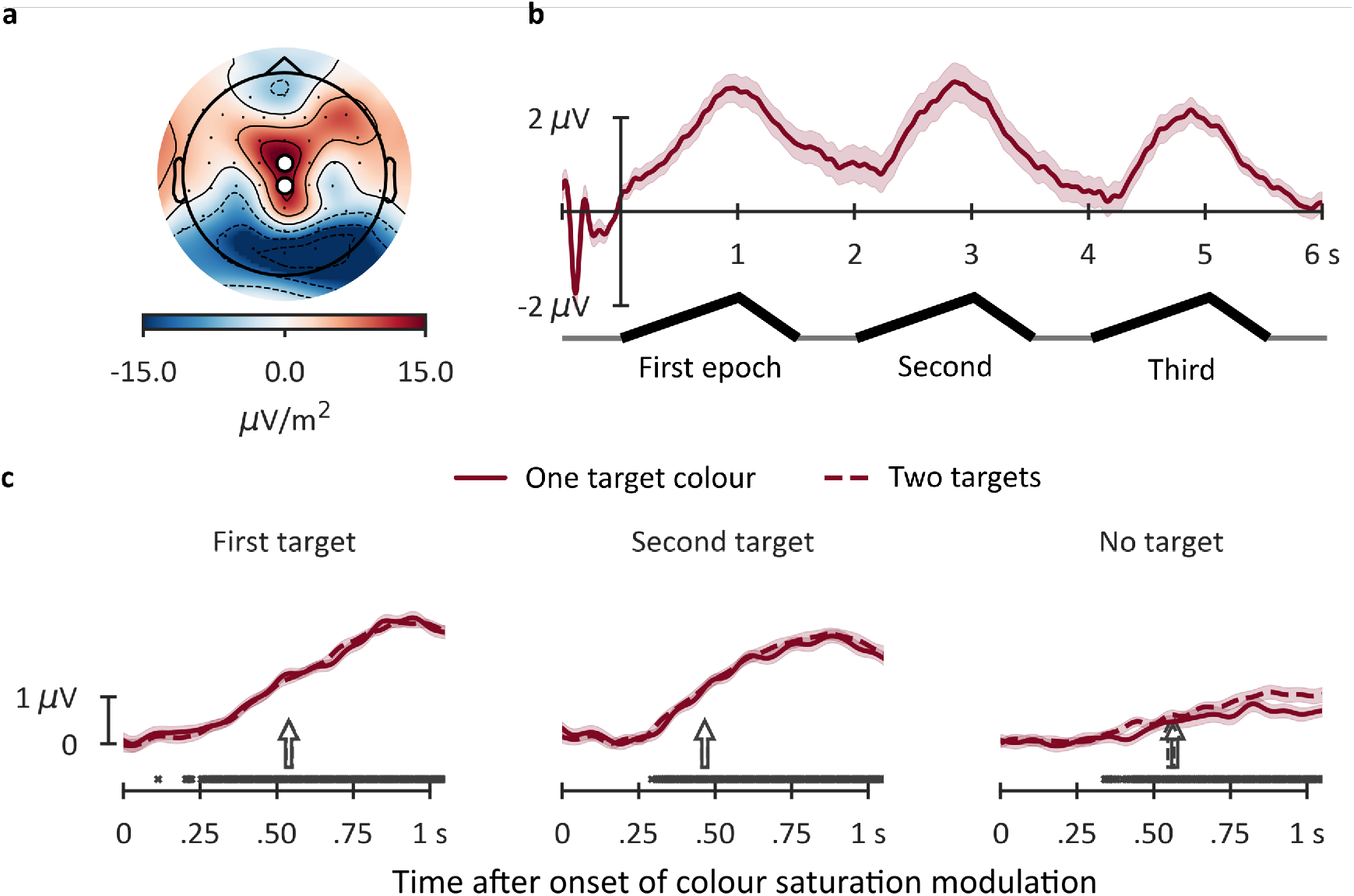
Topography and time course of the CPP component in Experiment 1. **a**, Surface-Laplacian-filtered EEG topography averaged across all conditions at times of peak colour saturation for all three epochs (1, 3, and 5 s after stimulus onset). The markers denote the electrode pair [CPz and Cz] used for the ERP analyses. **b**, Time-course of the CPP averaged across all experimental conditions (M ± 1 within-participants SEM; Morey, 2008) for the total trial duration. **c**, CPP time-course during the gradual increase in colour saturation, shown separately per epoch and experimental condition. Symbols above the x-axis denote time samples, averaged across experimental conditions, for which the voltage was significantly different from 0 at p_FDR_ < .05. Full-line and dashed-line arrows indicate onset latencies in the one and two target colours conditions, respectively. For the purpose of data presentation, all traces were low-pass filtered at 10 Hz using a 4^th^ order Butterworth IIR filter. Data analyses were conducted using unfiltered data.

The whole-trial epochs were then divided into three segments relative to the onset of colour saturation modulation, and baseline-corrected relative to the average of all three epochs over a window of 100 ms prior to onset. Thus, all three epochs had the same baseline, permitting comparisons between them. The epochs were then sorted according to experimental conditions (first target, second target, no target; one-and two-target colours) and averaged across trials (Fig. 2c). As we observed no effects of target colour repetitions vs. changes (all F_c_ < 1), all subsequent analyses were conducted using the average of these two trial types.

Overall, there was a statistically significant positive deflection in all experimental conditions (Fig. 2c). The onset latency (522 ms) and the CPP slope (3.31 μV/s) were similar across experimental conditions (all F_c_ < 1), suggesting that the rate of evidence accumulation did not differ between conditions. The peak CPP amplitude, however, *decreased* between epochs (F_c(2,64)_ = 46.41, p < .001), from the first target epoch to the second and then to the no-target epoch (2.74, 2.43 and 1.06 μV, respectively, all t_c(32)_ ≥ 1.93, p ≤ .031). Finally, the number of target colours (one vs two) had no significant effect on the peak CPP amplitude, either as a main effect (F_c_ < 1) or in combination with epoch (F_c(2,64)_ = 1.44, p = .244).

Taken together, the results of Experiment 1 revealed strong effects of selective attention on decision-making. Behaviourally, target signals were associated with significantly larger decision weights than distractors. At the neural level, attention had a robust effect on the peak CPP amplitude, with higher peaks for target-present epochs than for no-target epochs. Importantly, however, neither the CPP onset nor the slope differed between target-present and no-target epochs, suggesting that both task-relevant and task-irrelevant inputs triggered a process related to decision making. There was a robust and significant influence of distractor events on behaviour, suggesting that attention attenuates but does not fully suppress irrelevant inputs during integrated decision making. Significant distractor weights and the CPP ERP in the no-target epoch are consistent with the hypothesis that both target and distractor signals are integrated into a single decision variable.

In Experiment 1, we used an increase in colour saturation to gradually reveal to participants the task-relevance of the overlapping motion stimuli, and to render the coherent motion signals discernible. While this design was clearly effective for measuring the impact of distractors on decision making, it is possible that attentional filtering of distractors might have been more successful if participants had been given an opportunity to fully select the task-relevant stimuli (i.e., dot fields in the cued target colour) prior to the onset of coherent motion events. To test this hypothesis, in Experiment 2 the trial epochs were structured so that the colours of the dot stimuli reached full saturation prior to the onset of coherent motion events, thus effectively separating selection-related and response-related signals in time.

## Experiment 2

### Materials and Methods

#### Participants

36 healthy volunteers (20 females, average age 22.4 ± 4.4 years) participated in the study. All had normal or corrected-to-normal visual acuity and normal colour vision. Based on the same criteria as in Experiment 1, five participants were identified as outliers and removed from further analyses, leaving 31 participants in total. All other details were the same as in Experiment 1.

#### Stimuli, task and procedure

In Experiment 2 (Fig. 3a), the colour saturation increased rapidly (.25 s) from grey to full saturation, and remained at maximum saturation for 1 s. Coherent motion events were presented for .5 s during the maximum saturation period, and at variable times (.50–.75 s asynchrony relative to the onset of colour modulation).

To measure responses to targets and distractors independently, the dot fields corresponding to target and distractor signals flickered (ON-OFF) concurrently at different frequencies (12 and 18 Hz). The pairing between a given frequency and stimulus type (target vs. distractor) was counterbalanced across trials. The introduction of these flickering displays yielded distinct, time-resolved steady-state evoked potentials (SSVEPs, see Norcia et al., 2015), which enabled us to read out the strength of target and distractor representations on each trial. Previous research has shown that attention modulates the SSVEP amplitude with stronger responses to task-relevant than irrelevant stimuli (Andersen & Müller, 2010; O’Connell et al., 2012; Painter, Dux, Travis, & Mattingley, 2014) suggesting that the SSVEP amplitude is closely related to attentional selection dynamics.

**Figure 3.**
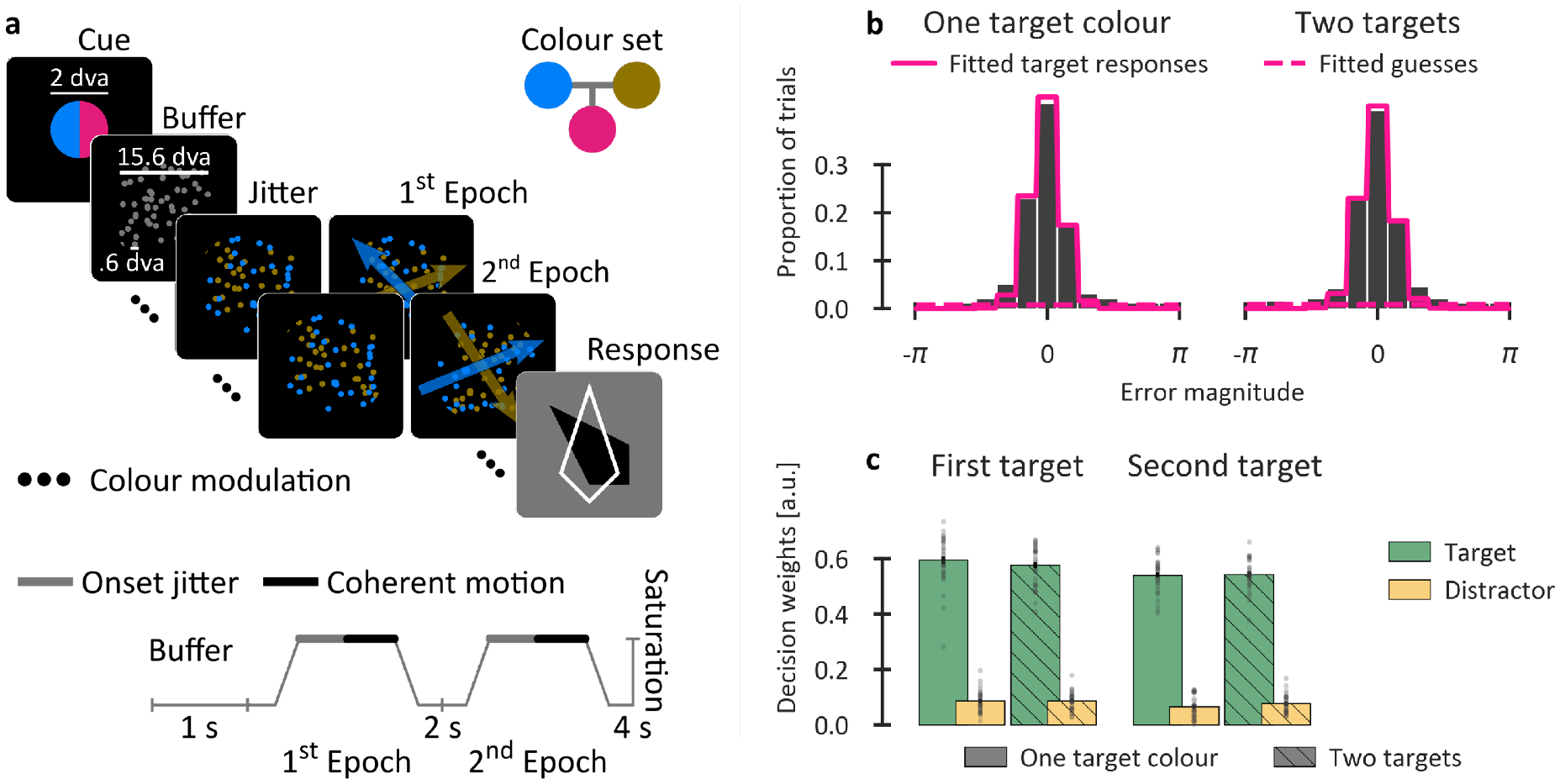
Schematic of the stimuli, task and the behavioural results in Experiment 2. **a,** Example stimulus sequence from a typical trial (above) together with the timing of colour saturation modulation and coherent-motion onsets (below). Only target-present epochs were presented, and the increase in colour saturation and onset of coherent motion were separated in time. Dot patches were frequency-tagged at 12 and 18 Hz (counterbalanced across target and distractor stimuli). **b**, Group-level histograms (bars) of the error magnitude together with the best-fitting mixture-distribution models (pink lines). **c**, Average decision weights showing the contributions of each presented motion direction to the decision. Grey dots denote individual participants. Conventions as in Figure 1.

To optimise the display for recording SSVEPs, the field size, dot size and dot number were doubled in Experiment 2 relative to Experiment 1. The dot-motion was generated at 2.5°/s, with an infinite dot life. Three distinctive colours (maximum saturation 75, minimum colour separation 90°) were selected; two colours served as targets, and the remaining one served as a distractor. The distractor and target colours were counterbalanced across participants. As the SSVEP analyses permitted independent tracking of target and distractor signals, only two epochs of dots were shown in Experiment 2, and both epochs within a trial contained a target-motion patch (i.e., there was no longer a “no target” epoch). Participants completed 64 practice trials followed by six blocks of 64 trials (384 trials in total). All other details were the same as in Experiment 1.

#### EEG analyses

In Experiment 2, time-resolved SSVEP power at tagged frequencies was computed using Morlet wavelets with fixed-width .5 s Gaussian taper, and normalized relative to the neighbouring frequencies. The mean power per epoch was computed during the 1 s of maximal colour saturation. For analyses of the target and distractor SSVEPs, the power per epoch was baseline-corrected relative to a .1 s interval around the onset of maximal colour saturation (.5 and 2.5 s, for the first and the second epochs, respectively). All other analyses were the same as in Experiment 1.

### Results

The behavioural results of Experiment 2 were qualitatively similar to those of Experiment 1, with a continuous unimodal distribution of error magnitudes centred at 0 (Fig. 3b). Initial analyses revealed no effect of repetitions vs. changes in the target colour (all ts < 1), so all subsequent analyses used the average of the two. Response precision (Fig. 3b) was high and comparable between one-and two-target colour trials (K/FWHM = 7.96/48° and 7.37/50°, respectively, F1,30 = 2.37, p = .134). The guessing rates were low and virtually identical for one-and two-target colour trials (ca. 12%, F < 1).

Analyses of decision weights revealed that both target and distractor signals contributed to the average motion-direction response in all conditions (all p < .001). The weights (Fig. 3c) for targets, however, were seven times higher than those for distractors (.56/.01 and .08/.01, respectively, F_1,30_ = 1,845, p < .001, 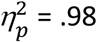). The serial order effect from Experiment 1 was replicated, with higher decision weights for the first epoch than for the second, both for targets (.58/.01 vs. .54/.01, t_30_ = 3.06, p = .004) and distractors (.09/.01 vs. .07/.01, t_30_ = 3.32, p = .002). Finally, there was no significant difference between one-and two-target colour trials, either as a main effect or in combination with the other factors (all F ≤ 1.35, all p ≥ .254).

We next characterised the time-course of the CPP. The ERP topography (Fig. 4a) in Experiment 2 was broadly consistent with that of Experiment 1, showing a central-medial positivity. The time-course of the CPP (Fig. 4b) throughout the trial closely followed the modulation of colour saturation, peaking sharply around .8 and 2.8 s and decaying quickly thereafter. There was also a second modulation, broadly coinciding with the coherent motion onsets (at around 1.5 and 3.5 s), which, due to the jittered timing of the motion onsets, rose and fell more slowly than the colour-evoked components.

**Figure 4.**
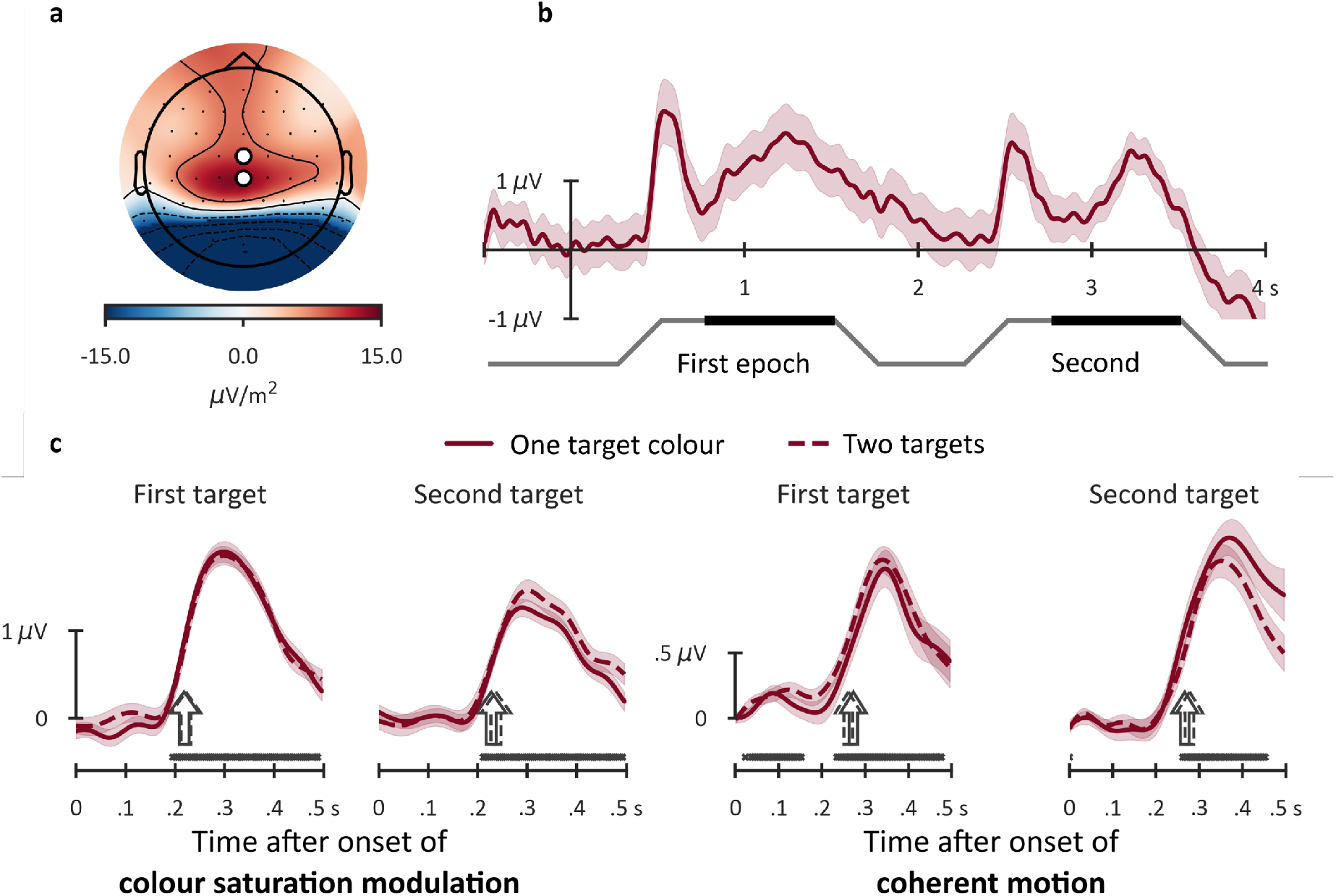
Topography and time course of the CPP component in Experiment 2. **a**, Surface-Laplacian-filtered EEG topography averaged across all conditions at the middle of peak colour saturation for both epochs. **b**, The CPP component averaged across all experimental conditions for the total trial duration. **c**, Time-course of the CPP time-locked to the onset of colour saturation (left panels), and to the onset of coherent motion (right panels), shown separately per epoch and experimental condition. Note the difference in scale between panels. Conventions as in Figure 2.

Analyses of the CPP (Fig. 4c) time-locked to the onset of colour modulation and coherent motion, respectively, revealed a significant positive deflection in both cases. Analyses of ***colour-locked CPPs*** for the first epoch relative to the second revealed a shorter onset latency (220 vs. 231 ms, F_c(1,30)_ = 5.04, p = .032), a marginally steeper slope (17.57 vs. 13.34 μV/s, F_c(1,30)_ = 3.23, p = .083) and a higher peak amplitude (1.95 and 1.42 μV, F_c(1,30)_ = 6.40, p = .017). In the second epoch, dividing attention between two target colours yielded a significantly higher peak amplitude for two-versus one-target colour trials (1.56 and 1.29 μV, respectively; t_c(1,30)_ = 2.30, p = .029) . Analyses of ***motion-locked CPPs***, by contrast, revealed no significant differences in onset latency (270 ms), slope (9.44 μV/s) or peak amplitude (1.27 μV) between experimental conditions (all F_c_ <= 1.44, all ps >= .240). Overall, the ERP analyses of Experiment 2 suggest that discerning the task-relevant colour patch (i.e., selection-relevant feature) and the coherent motion direction (i.e., response-relevant feature) both engaged decision-making processes to some degree. Further, as only the colour-locked CPP differed between the first and second epoch, it appears that the serial order effect observed in both Experiments 1 and 2 was more closely related to processing of colours rather than processing of the motion signal itself.

To independently characterise the brain’s response to target and distractor signals, we next analysed the amplitude of SSVEP responses elicited by the frequency-tagged motion stimuli. There was a strong (≈2 dB) frequency-specific response which peaked at the Oz electrode over the occipital pole (Fig. 5a,b). Time-frequency analyses revealed that the power of both target and distractor signals was significantly different from zero throughout the trial (all p_FDR-corrected_ < .05). Moreover, the power was similar for target and distractor patches prior to the increase in colour saturation in both the first and second epochs (Fig. 5c).

**Figure 5.**
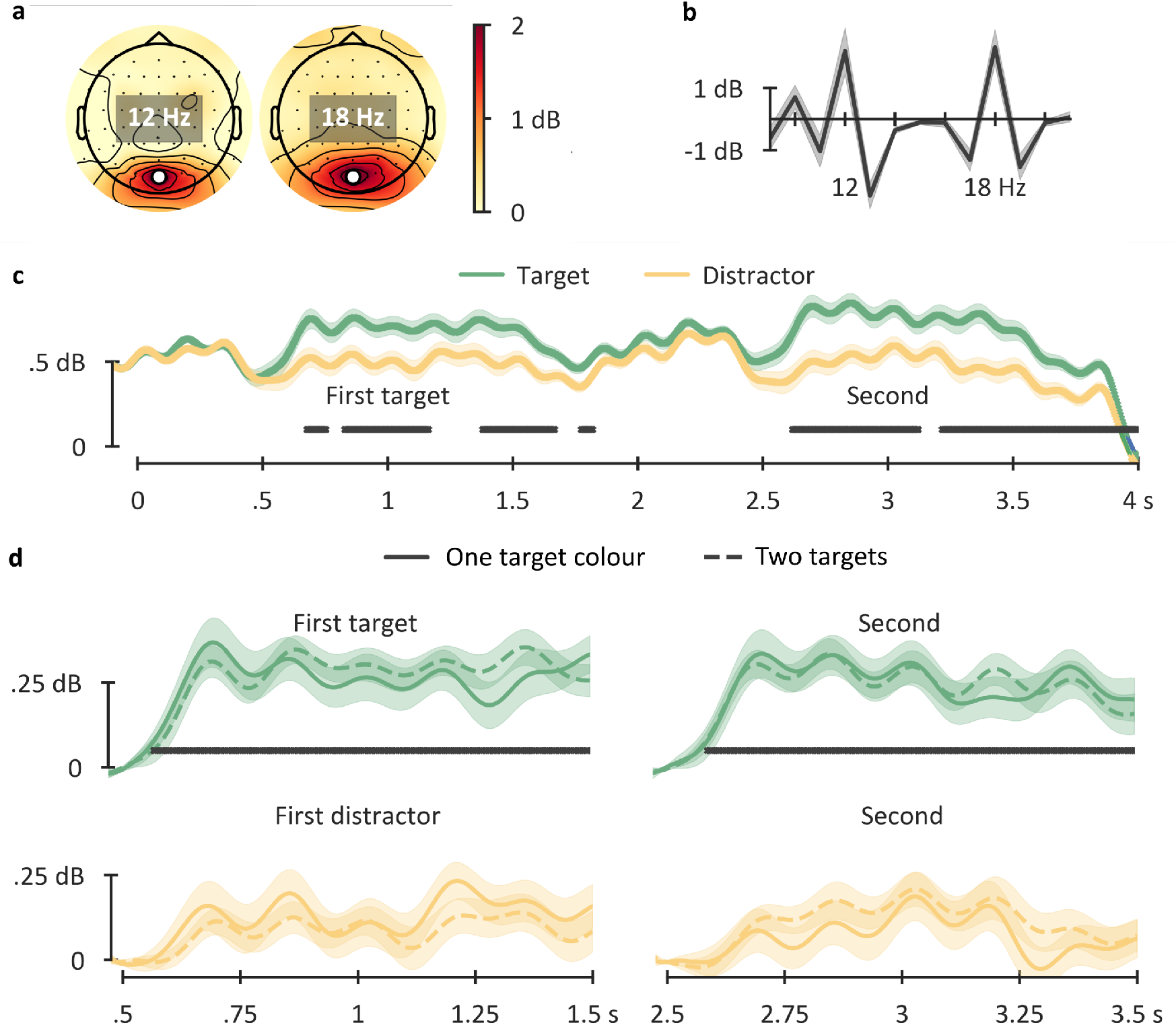
Time-course of target and distractor processing measured as SSVEP power in Experiment 2. **a**, Topography of the total power at 12 Hz and 18 Hz normalized relative to the neighbouring frequencies, computed using a fast-Fourier transform during periods of maximal colour saturation (.5–1.5 s, and 2.5–3.5 s). **b**, Normalized total SSVEP power (M ± 95% CI) at Oz electrode during the period of maximal colour saturation averaged across epochs and all experimental conditions. **c**, Time-resolved target-and distractor-evoked SSVEPs for the whole trial duration averaged across all experimental conditions (M ± 1 within-participants SEM). Black symbols above the x-axis denote times at which the difference between target and distractor power was significant at p_FDR-corrected_ < .05. **d**, Target-evoked (upper panels) and distractor-evoked (lower panels) SSVEPs per epoch and per experimental condition baseline-corrected relative to .1 s interval around the onset of maximum colour saturation (.5 and 2.5 s for the first and second epoch, respectively). Black symbols above the x-axis denote times at which the baseline-corrected power was significantly different from zero at p_FDR_ < .05.

As the colour saturation increased, there was a transient *decrease* in power for the two patches in both the first epoch (around .5 s) and the second (around 2.5 s). Whereas the power for target patches in both epochs recovered quickly (Fig. 5d, upper panels), the power for distractor patches remained unchanged throughout the epochs (Fig. 5d, lower panels), yielding a robust effect of task-relevance (average power: .24 and .10 dB for target and distractor patches, respectively, F_1,30_ = 11.64, p = .002, 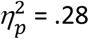). Finally, the SSVEP power was comparable across the first and second epochs, and across one-and two-target colours (all F ≤ 1.25, all p ≥ .272).

## Analyses of feature-specific brain responses

In a final analysis, we characterised the strength of feature-specific brain responses to coherent motion events in both Experiments 1 and 2 (Fig. 6a,b). Inspection of the motion tuning in Experiment 1 revealed a phasic response to both target and distractor motion 0–.5 s after the onset of coherent motion. During the following 1 s period, a feature-specific response developed gradually, peaking at around 1.5 s. Interestingly, throughout the epoch, motion tuning was comparable for the target and distractor motion directions. To test the effect of stimulus type (target vs distractor) on tuning strength, we conducted stepwise linear mixed effects modelling with participant as a random effect, separately for each time sample. In the first step, the model included only the average tuning strength per sample (the intercept). In the second step, the stimulus type (target vs distractor) was added as a fixed effect, and the increase in the goodness-of-fit between different models (Intercept only vs Intercept + Stimulus) was assessed using a log-likelihood ratio test. The resulting p-values for all time samples were corrected for multiple comparisons (FDR-correction). The modelling revealed overall significant tuning at around .25 s and 1.5 s (Fig. 3a, lower panel, the intercept). The effect of stimulus did not reach significance at any time point.

**Figure 6.**
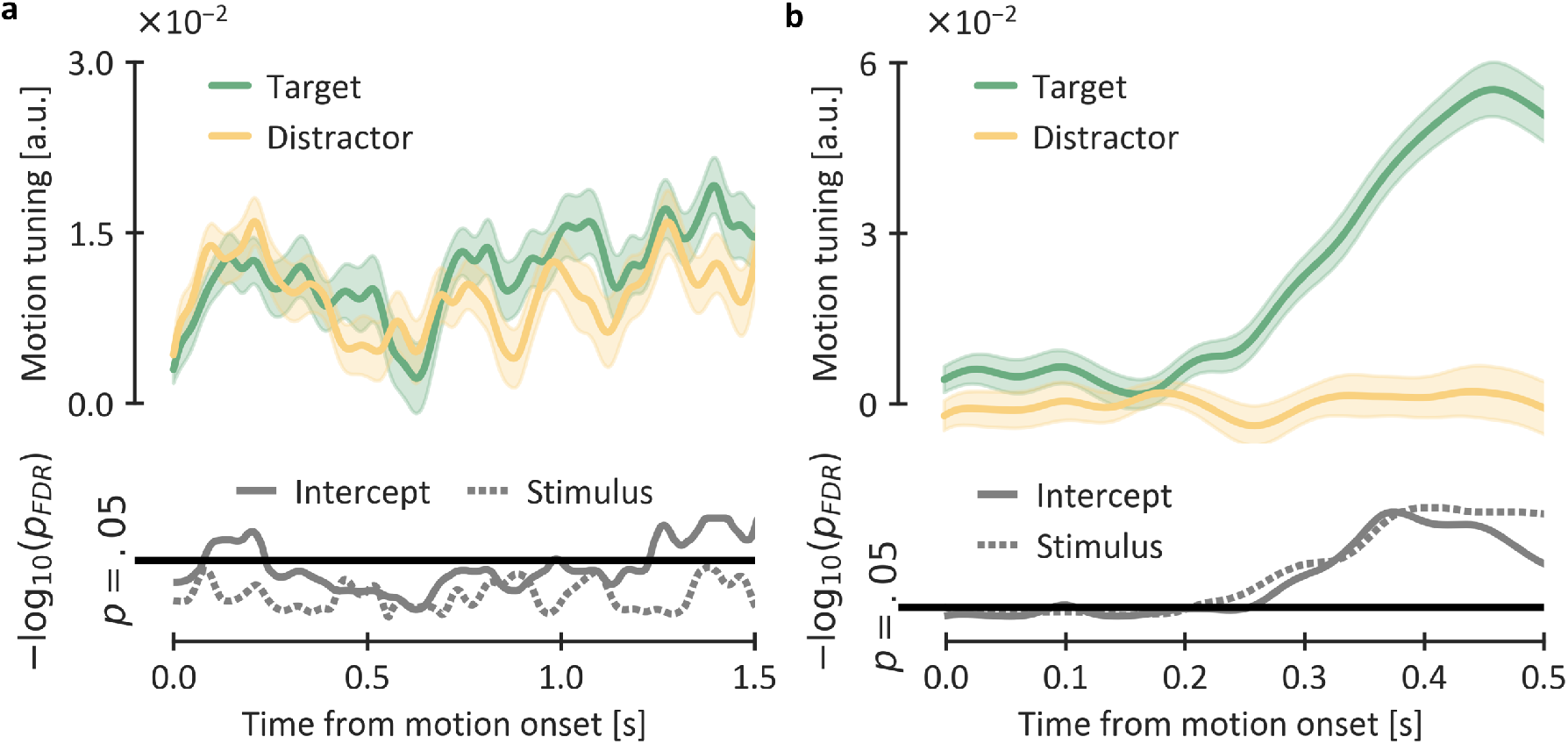
Time-course of feature-specific responses to coherent motion direction (i.e., motion tuning). **a**, Upper panel shows tuning strength (M ± 1 within-participants SEM) averaged across one-and two-target colours, and first and second target epochs, in Experiment 1. Higher values indicate stronger tuning. Lower panel shows FDR-corrected p-values (−log10-transformed) for the overall tuning strength (Intercept) and the effect of stimulus (Target vs Distractor) per time-sample. The horizontal line denotes .05 cut-off. **b**, Tuning strength in Experiment 2. Conventions as in panel a. Note the difference in x and y scales between panels a and b.

Inspection of motion tuning in Experiment 2 revealed strong tuning to target motion, starting from around .2 s after the onset of coherent motion. By contrast, tuning to distractor motion was negligible throughout the epoch. These observations were confirmed by a stepwise LME model which revealed both strong motion tuning overall (the intercept) and a strong effect of stimulus (target vs distractor). Relative to the tuning analyses conducted in Experiment 1, tuning to target motion direction in Experiment 2 was roughly four times stronger, consistent with the paradigm differences between experiments. More specifically, whereas in Experiment 1 participants had to concurrently select the task-relevant dot field and discern its motion direction, in Experiment 2 they could first select the target field and then dedicate themselves to processing motion direction.

Overall, the results of the motion tuning analyses in Experiments 1 and 2 are broadly consistent with the behavioural decision weights in the two experiments. In Experiment 1, the behavioural target weights were four times stronger than the distractor weights, whereas in Experiment 2 target weights were around seven times stronger. Similarly, the tuning strength was overall numerically lower in Experiment 1 than in Experiment 2. These results suggest that providing additional time for attentional selection (as in Experiment 2) increases the precision with which task-relevant features are represented and decreases the influence of concurrently presented task-irrelevant features.

## Discussion

The goal of the present study was to characterise the behavioural and neural processes associated with making integrated perceptual decisions. Unlike simple perceptual decision-making (Forstmann et al., 2016; Gold & Shadlen, 2007; Ratcliff et al., 2016), which requires judgments on a single, task-relevant stimulus, our decision-making task involved selective processing of target signals in the presence of distractors, and required integration of sensory evidence over an extended time period within a trial. The two main questions we sought to address in this study were: (i) whether attentional selection and decision making take place in parallel or sequentially, and (ii) whether task-relevant and task-irrelevant sensory inputs are accumulated into a single decision variable.

To characterise the role of selective attention in decision making, we used mixture distribution modelling and regression analyses. To characterise the neural correlates of decision-making, we used a widely accepted correlate of decision making in humans, the centro-parietal positivity (CPP) component (Hanks & Summerfield, 2017; O’Connell et al., 2018; Spitzer et al., 2017). To monitor processing of different, task-relevant and task-irrelevant sensory inputs, in Experiment 2 we flickered stimuli at different frequencies and measured SSVEPs to targets and distractors. Finally, to characterise feature-specific brain responses to coherent motion direction, we used a multivariate analysis approach that was successfully used in the past to characterise feature-specific responses in a variety of tasks (Ede et al., 2018; Wolff et al., 2017).

In Experiment 1, there was a measurable CPP in no-target epochs, coupled with a robust and significant effect of distractor motion signals on behaviour. Taken together, these results suggest that participants were unable to suppress distractor signals from being accumulated into the decision variable. Importantly, both the peak CPP amplitude and the decision weights in the no-target epochs were lower than the CPP amplitude and target weights in target-present epochs, suggesting that selective attention modulated the contribution of distractor motion to decision making. Consistent with Experiment 1, the neural responses to distractors in Experiment 2, while significantly smaller than those to targets, were not completely suppressed and remained stable over the period during which target and distractor patches were discernible within the trial. There was also a statistically significant distractor effect on responses, albeit smaller than the distractor effect in Experiment 1. Finally, we also observed qualitative differences between Experiments 1 and 2. Specifically, the behavioural target weights were numerically higher and the distractor weights numerically lower in Experiment 2 than Experiment 1. Similarly, the motion tuning analyses revealed a significant difference in tuning to target and distractor motion directions in Experiment 2, whereas there was no such difference in Experiment 1.

Taken together, the results of the two experiments suggest that: (i) selective attention unfolds in parallel with evidence accumulation, and that (ii) both target and distractor signals are accumulated into a single decision variable. As attentional selectivity increases, distractor inputs are attenuated. Importantly, our findings demonstrate that attention does not completely suppress the accumulation of task-irrelevant sensory input. The differences between Experiments 1 and 2 can be parsimoniously explained by differences in the experimental paradigm: in Experiment 1, coherent motion onset within each trial coincided with modulation of colour saturation, whereas in Experiment 2 motion onset followed modulation of colour saturation. Assuming that attentional selectivity at the onset of coherent motion differed between experiments accounts for the discrepancies between decision weights and motion tuning in a straightforward way.

Overall, the results of the present study are consistent with the notion that sensory evidence accumulates in parallel (rather than sequentially) with an increase in selectivity for task-relevant features. Computational modelling of response speed and accuracy in the classic flanker task – in which target and distractor stimuli are presented concurrently at different spatial locations (e.g., Eriksen, 1995) – has shown that similar principles apply to selectivity for task-relevant locations (White et al., 2011). Of note, in a recent study Wyart et al. (2015) had participants report the average orientation of one of two spatially separated streams of achromatic gratings. Unlike the present study, decision weights for the spatially separated distractors in their study were statistically indistinguishable from zero. While differences in experimental tasks and analytical approaches prevent direct comparisons between our study and that of Wyart et al. (2015), the two sets of findings suggest that space-and feature-based filtering of sensory inputs during evidence accumulation may exhibit different properties. Specifically, sensory evidence accumulation might be restricted to a finite region of space so that under conditions in which signals are sufficiently separated they engage independent accumulators.

Interestingly, across both the current experiments, having participants divide attention between two potential target colours yielded no behavioural cost relative to having them focus on a single target colour. In addition, there was no evidence of a cost for switching between target colours across two epochs, compared with trials in which the two colours repeated. Since in the present study the target colour patches were rendered increasingly visible over a prolonged time interval and the responses were not speeded, it is possible that our paradigm was insensitive to potential behavioural costs of dividing attention (but see, e.g., Eimer and Grubert, 2014; Grubert et al., 2017). On the other hand, the neural correlates of decision making, as characterised by the onset latency, slope, and peak CPP amplitude, were consistent with the behavioural findings: the CPP time-traces were virtually identical for one-and two-target colour trials in both experiments. As the specifics of our experimental design prevent any strong conclusions, these findings call for further investigation.

In both experiments, we also found a significant serial order bias with higher decision weights for the first target epoch relative to the second. In Experiment 1, higher target weights were accompanied by lower distractor weights, whereas in Experiment 2 the weights for both target and distractor signals were higher in the first than the second epoch. This primacy effect was not expected and, in fact, studies using similar paradigms have typically reported either no order effects or even the opposite, recency effects (Cheadle et al., 2014; Wyart, de Gardelle, Scholl, & Summerfield, 2012; Wyart et al., 2015). Mirroring the behavioural primacy effects, the colour-locked peak CPP amplitude in Experiment 2 was higher for the first than the second epoch. By contrast, analyses of the motion-locked CPP revealed no order effects. These results suggest that the primacy effects might be related to processing colours rather than motion. Since previous studies did not modulate attention-related signals in the way we did here, the primacy effects we observed are not necessarily inconsistent with the literature. Further studies will be necessary to characterise this effect in more detail.

The motion averaging task we have introduced here has potential as a tool for investigating the mechanisms underlying more complex, integrated decision-making. Dynamic weighting of individual decisions to inform a single, integrated decision is ubiquitous in daily life: for example, when deciding whether or not to cross a busy road. In our task, individual motion signals are uncorrelated and, considered in isolation, are not predictive of the final decision. Hence, the averaging task permits independent parameterisation of both *simple* and *integrative* decision-making processes in a straightforward and well characterised motion discrimination task. Using neuroimaging methods with high spatial resolution such as fMRI, future studies could identify the brain areas involved in simple decision-making, on the one hand, and integrative decision-making, on the other. Using this task to characterise decision-making across the lifespan or in clinical groups could also shed light on the development of integrative decision-making abilities which are a hallmark of adaptive, intelligent behaviour.

## Acknowledgments

This work was supported by the Australian Research Council (ARC) Centre of Excellence for Integrative Brain Function (ARC Centre Grant CE140100007). JBM was supported by an ARC Australian Laureate Fellowship (FL110100103). The funders had no role in study design, data collection and analysis, decision to publish, or preparation of the manuscript.

## Notes

*Conflict of interest:* The authors declare no competing financial interests.

#### Summary of Updates

Multivariate feature decoding analyses were added to Results.

https://doi.org/10.14264/uql.2019.7

